# Cocaine Self-Administration and Time-dependent Decreases in Prelimbic Activity

**DOI:** 10.1101/255455

**Authors:** Torry S. Dennis, Thomas C. Jhou, Jacqueline F. McGinty

## Abstract

Cocaine self-administration causes dephosphorylation of proteins in prelimbic (PL) cortex that mediate excitation-transcription coupling, suggesting that cocaine causes decreased activity in PL neurons. Thus, we used *in vivo* single-unit extracellular recordings in awake, behaving rats to measure activity in PL neurons before, during, and after cocaine self-administration on the first and last session day (range 12-14 days). On the last day, cocaine suppressed 52% of recorded neurons in the PL cortex when compared to a 20 min baseline, significantly more than the 23% of neurons suppressed on the first day of cocaine. There was no change in the percentage of neurons that were excited on the first vs. the last day of self-administration (14% vs. 13%, respectively). To determine whether the tonic inhibitory shift was induced by the behavior or by the drug itself, rats received a final session in which cocaine availability was delayed for the first 30 min or in which cocaine was administered non-contingently in the absence of levers or cues in a subset of rats. These manipulations indicated that cocaine was both necessary and sufficient to induce a downshift in PL neuronal activity. However, this activity decrease was not observed in rats that received only non-contingent cocaine for 12-14 days, indicating that contingency during self-administration training contributes to the cocaine-induced tonic downshift in PL activity. These data suggest that the session-by-session decrease in PL neuronal activity induced by cocaine is a reflection of learned drug-seeking behavior and, hence, reducing this inhibition may lessen cocaine addiction.

## INTRODUCTION

This study examined the effect of repeated cocaine self-administration (SA) on neuronal firing in the prelimbic (PL) cortex before, during, and immediately after SA. The PL cortex has an established involvement in drug-related behaviors: PL inactivation significantly reduces cocaine SA (Di Pietro *et al*, 2006; Gutman *et al*, 2016) whereas PL inactivation immediately prior to reinstatement suppresses context (Fuchs *et al*, 2005), cue (McLaughlin and See, 2003; Stefanik *et al*, 2016), cocaine-primed (Di Pietro *et al*, 2006; McFarland and Kalivas, 2001), and stress-induced (Capriles *et al*, 2003; McFarland *et al*, 2004) drug seeking. Corresponding changes take place in the PL cortex with continued cocaine use that contribute to relapse. Previous work from our laboratory has identified profound dephosphorylation of proteins that mediate excitation-transcription coupling (pERK, pCREB, pGluN2A, and pGluN2B) in the PL cortex within two hr after the end of cocaine SA (Go *et al*, 2016; Whitfield *et al*, 2011). A single infusion of brain-derived neurotrophic factor (BDNF) into the PL cortex immediately after the end of SA reverses these changes (Go *et al*, 2016; Whitfield *et al*, 2011) and normalizes cocaine-induced dysregulation of glutamate levels in the nucleus accumbens (Berglind *et al*, 2009) that are associated with relapse (McFarland *et al*, 2003). Importantly this same treatment suppresses subsequent context, cue, and cocaine-primed reinstatement (Berglind *et al*, 2007) in a Trk- and ERK-dependent manner (Whitfield *et al*, 2011). These studies highlight the importance of characterizing neuroadaptations in PL cortex during and immediately after repeated cocaine SA.

Additional support for the importance of PL cortex in drug-related behaviors comes from the use of *in vivo* electrophysiological recordings in awake, behaving animals. This approach allows for the analysis of complex patterns of activity and their evolution across time in individual animals. For example, cocaine SA has been shown to decrease PL cortical baseline activity over multiple sessions (Sun and Rebec, 2006). At the same time, this technique also provides high temporal resolution and allows for the direct comparison of physiological measurements with specific behaviors or experiences. For example, measurements in the PL cortex have revealed the presence of anticipatory changes (both increases and decreases) in neuronal activity directly preceding the lever press to receive cocaine (Chang *et al*, 2000; West *et al*, 2014) and both increases (Sun and Rebec, 2006) and decreases (Chang *et al*, 1997; 1998) in activity directly following the cocaine infusion. Larger trends in activity can also be explored across an entire behavioral session. Of particular interest is a study showing that cocaine SA induces tonic decreases in the majority of medial prefrontal neurons (mostly but not exclusive in the PL cortex) in rats undergoing concurrent heroin and cocaine self-administration (Chang *et al*, 1998). While these studies have expanded our understanding of the role of the PL cortex in drug-taking behaviors, there are still unanswered questions. For example, do the observed changes in activity develop with repeated cocaine use or are they present on the first day of SA? Additionally, what is the underlying behavioral or pharmacological etiology of these changes in activity? Lastly, is there any relationship between session-long tonic activity changes induced by cocaine SA and phasic firing surrounding discrete drug-seeking behaviors? The current set of experiments was designed specifically to answer these questions.

## METHODS

### Subjects

Adult male Sprague Dawley rats (n=34) were purchased from Charles River Laboratories (Wilmington, MA) at 276-300g. Rats were individually housed and kept in a temperature and humidity controlled facility on a reversed 12-hr light/dark cycle (lights off at 7:00am). Food and water were supplied *ad libitum* upon arrival and through surgical recovery. When behavioral procedures were initiated, rats were restricted to 20g of rat chow daily. All experimental protocols were approved by the Institutional Animal Care and Use Committee (IACUC) of the Medical University of South Carolina and were performed in accordance with the National Institutes of Health Guide for the Care and Use of Laboratory Animals (8^th^ ed., 2011).

### Surgery

Rats were allowed to acclimate for at least three days before any surgical procedures were performed. On surgery day, rats were anesthetized with ketamine (66 mg/kg, i.p.) and xylazine (1.33 mg/kg, i.p.). Ketorolac (2 mg/kg, i.p.) was administered for analgesia. Silastic tubing was implanted into the right jugular vein and connected to a chronically indwelling catheter pedestal (Plastics One, Roanoke, VA), externalized 1 cm posterior to the shoulder blade through a small incision in the skin. Catheter patency was sustained through daily flushing of a taurolidine/citrate solution (0.05 ml, i.v.; Access Technologies, Skokie, IL). Immediately after the catheter surgery was completed, anesthetized rats were transferred to a stereotaxic frame (Kopf Instruments, Tujunga, CA) for electrode implantation. A custom made 16-wire (25 µm Formvar-Coated Nichrome; A-M Systems, Sequim, WA) drivable electrode bundle was housed within a 31-gauge polyimide tube and unilaterally implanted into either the right or left PL cortex (coordinates A/P: +3.0, M/L ±0.6, D/V 1.8 from dura-Paxinos and Watson, 2005). A grounding pin was implanted into a separate area of the posterior cortex. The electrode bundle was fixed to the skull using acrylic cement with 5-6 steel skull screws for additional anchoring. Electrode wires were driven further into the PL cortex after surgery but before the initiation of behavioral procedures. Rats received postoperative care for 5 days following surgery.

### Behavioral Procedures

#### Cocaine Self-Administration

Rats were trained to self-administer cocaine on an FR 1 schedule of reinforcement. Training occurred in standard operant chambers containing two retractable levers, a house light, a tone generator, and a signal light above each lever (Med Associates, Inc). The operant chambers were housed within sound attenuating boxes. An active lever press resulted in an i.v. cocaine infusion (0.2 mg/50 µl/infusion) and a 5 sec tone-light cue complex, followed by a 20 sec timeout period in which lever presses had no programmed consequence. Presses on the inactive lever were recorded but had no consequence. Within each session, rats were given a 20-min baseline period in which the house light was illuminated but no levers were present. After 20 min, levers were extended and rats were allowed to self-administer cocaine for 2 hr. After 2 hr of SA, levers were retracted, and rats remained in the chamber with continuous electrophysiological recording for an additional 2.5 hr. SA criterion was &≥ 10 infusions/day during 12-14 days. After the SA paradigm was completed, a subset of rats (n=9) underwent a session on the next day (Experiment 2a) in which the availability of cocaine was delayed for the first 30 min of the SA session (levers were available and cues were paired with lever presses during the delay). After two additional cocaine SA sessions (Experiment 2b), these rats were non-contingently administered cocaine randomly in the operant chamber for 2 hr in the absence of levers and cues. For the latter procedure, the number of infusions was determined by averaging infusions for the last three days of SA for each individual rat.

#### Yoked Cocaine

A separate cohort of rats received yoked cocaine administration throughout the entire paradigm (12-14 days) in Experiment 3. The infusion schedule for these rats was yoked to the infusion schedule of previously self-administering rats. Cocaine infusions were always paired with a 5-sec light-tone cue complex. Responses on either of the two levers yielded no programmed consequence. Within each session, rats underwent a 20 min baseline period, after which levers were extended. Rats then received two hr of yoked cocaine administration (0.2 mg/50 µl/infusion). Levers were then withdrawn and rats remained in the chamber for an additional 2.5 hr. Electrophysiological recordings were taken on the first and last day of yoked cocaine administration.

### Electrophysiological Recording & Analysis

Electrophysiological signals were received through a 16-channel headstage and preamplifier (Neurosys LLC, Charleston, SC). Analog signals from the preamplifier were converted to digital signals using a PCI A/D converter (National Instruments, Austin, TX). Signals were initially processed/recorded using Neurosys Acquisition Software. Electrophysiological data was further processed using Neurosys Offline Spike Sorter software, NeuroExplorer (Nex technologies), and Matlab (Mathworks). Isolated neurons were accepted if a refractory period was evident, as indicated by a large central notch in an autocorrelogram. The “first day” of electrophysiological recording was defined as the first day in which animals received at least 5 infusions of cocaine.

### Experimental Design

Figure 1 shows the timeline for all experimental procedures. Experiment 1 was conducted to characterize changes in neuronal activity that occur before, during, and immediately after cocaine SA. Electrophysiological recordings were taken on both the first and last day (the last day ranged from 12-14 days) of SA to examine changes in patterns of neuronal activity that occur not only within each session, but also with continued cocaine use over time.

**Figure 1:**
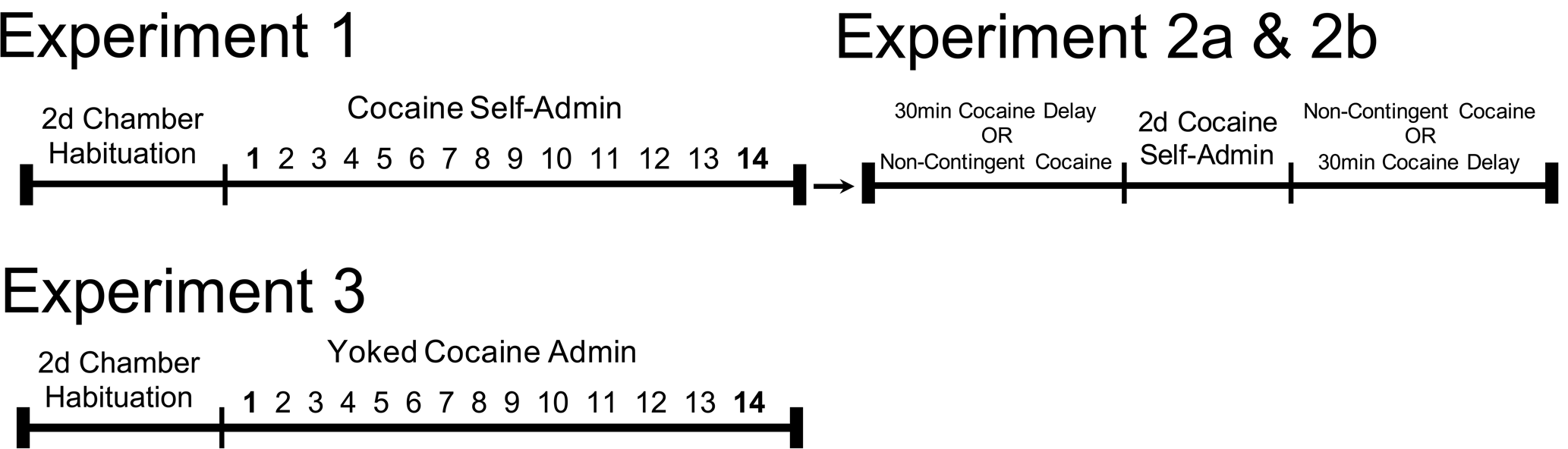
Timeline for Experiments 1-3.

The purpose of Experiment 2a was to determine if cocaine was necessary to elicit transient tonic changes in neuronal activity in animals with a history of cocaine self-administration. The purpose of Experiment 2b was to determine if cocaine alone was sufficient to elicit transient tonic shifts in neuronal activity in animals with a history of cocaine self-administration. The order of experiment 2a and 2b was counterbalanced, with two days of standard self-administration in between. The purpose of Experiment 3 was to determine whether transient tonic shifts in activity observed from early to late self-administration in Experiment 1 were due to the pharmacological effect of cocaine administered over 12-14 days. To determine this, a separate cohort of animals was yoked to the infusion schedule of previously self-administering rats and non-contingently administered cocaine for 12-14 days. As in Experiment 1, electrophysiological recordings were taken on the first and last day.

### Histology

After the behavioral paradigm was complete, rats were anesthetized and perfused with 10% buffered formalin / 2% potassium ferrocyanide. After the perfusion, but before the brain was extracted, a 200 µA current was passed through each electrode wire for 5 sec. The iron deposits from the electrode reacted with the potassium ferrocyanide to form a Prussian blue stain at the tip of the electrode. Brains were sliced at 40 µm on a freezing microtome. Tissue was mounted on a slide and Nissl-stained. Only data from electrode placements that were fully contained within the PL cortex (Paxinos and Watson, 2005) were used in the final analyses (Figure 2). Four animals were excluded for missed placement.

**Figure 2.**
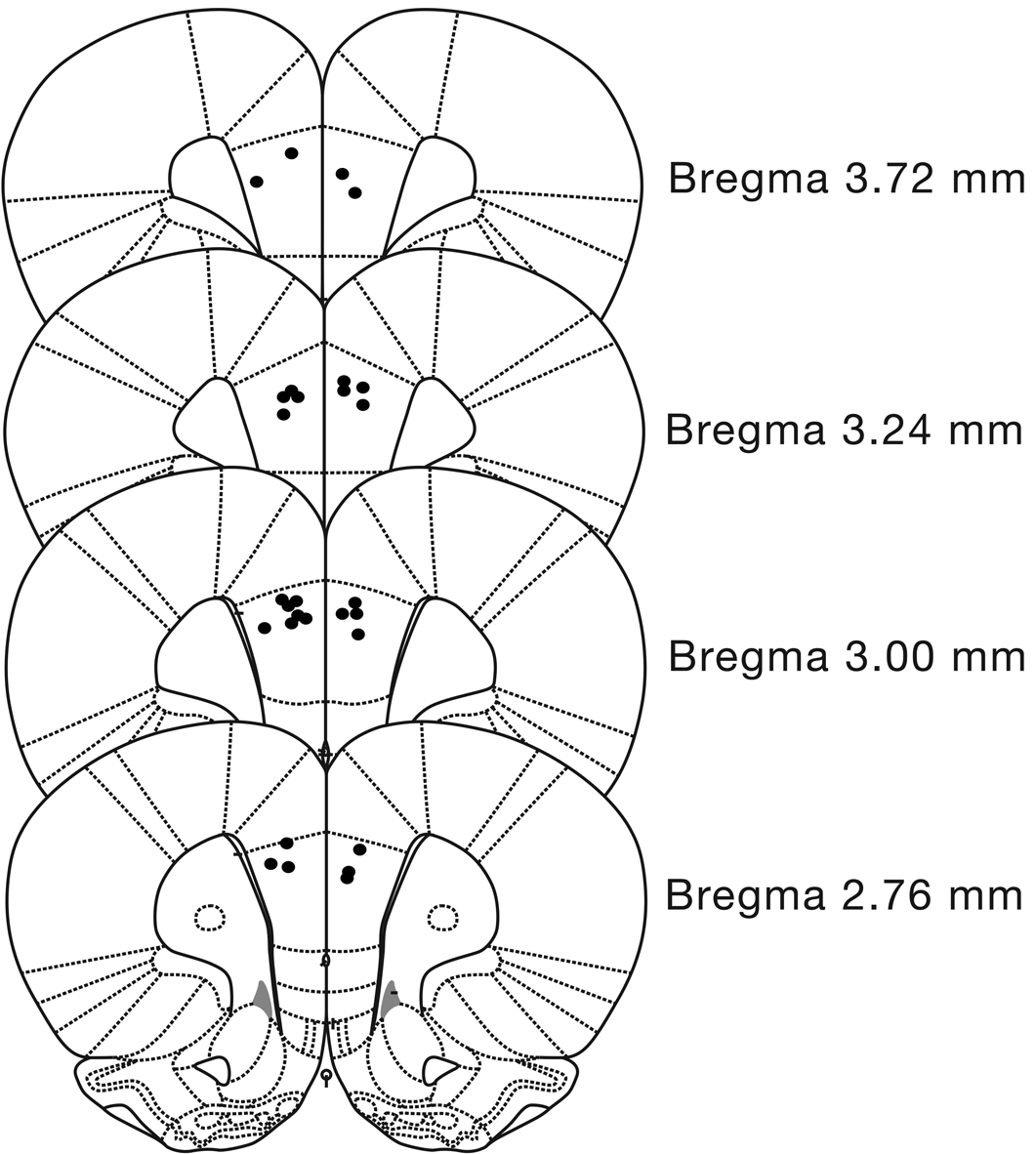
Graphical representation of histological placements from Experiments 1-3. Each dot represents the tip of the implanted electrode bundle aimed at the PL cortex. Data from 4 rats with placement outside the PL cortex were excluded from analysis.

### Statistical Analysis

To identify overall changes in the firing of an individual neuron within a given session, spike rates were normalized by dividing average firing during self-administration and early withdrawal by the average firing rate for the 20-min baseline of that session. This provided a “fold change” from baseline firing for each day of recording. A criterion of two standard deviations from the normalized average firing rate was used to screen outliers from the dataset. Cells were then divided into one of three subgroups, depending on the extent of deviation from baseline to self-administration (> 25% decrease, >25% increase, & all cells that fell in between). One sampled t-tests, with Bonferroni correction for multiple comparisons, were used to compare firing rates during self-administration and early withdrawal to the baseline for each subgroup. Because the normalized baseline equals 1, the theoretical number of 1 was used as the comparison in the one sampled t-tests. To identify changes in the distribution of neurons that are excited/suppressed during self-administration from one session to another, a chi-square was used. Firing rates were also recorded 10 sec before and 10 sec after the lever press that resulted in a cocaine infusion. These data were grouped into the three subcategories, as before, depending on the extent of deviation in firing from the 20-min baseline to self-administration (> 25% decrease, >25% increase, & all cells that fell in between). Rates of firing were normalized by dividing each data point by the average firing rate of each neuron from -15 to -10 sec prior to the lever press. This provided a “fold change” for activity in this time range. For statistical analysis, electrophysiological firing was grouped into 2 sec bins and one sampled t-tests with Bonferroni correction were conducted against the theoretical number of 1 for each subgroup. The criterion for statistically significant differences for both t-tests and chi-square tests was p<0.05.

## RESULTS

### Histology

The electrode placements in the PL cortex of rats in all experiments are depicted in Fig 2.

### Experiment 1: Cocaine SA decreases overall activity in the majority of recorded neurons in the PL cortex after 12-14 days of cocaine self-administration

Qualitative changes in the pattern of firing and quantitative changes in the distribution of cells that were excited/inhibited in the PL cortex were analyzed before, during, and after cocaine SA on the first and last day. The percentage of cells that were inhibited during the 2-hr SA period significantly increased from 23% on the first day (Figure 3A) to 52% on the last day (Figure 3B) (Chi-Square: X^2^ (2, N=265) = 25.19, p<0.0001). On the first day of SA, cells that were inhibited during SA took 80 min on average to return to baseline rates of firing (Figure 3A). By the last day of SA, inhibited neurons returned to baseline rates of firing within the first 20 min of withdrawal (Figure 3B). Firing rates were also calculated for time windows ±10 seconds before and after each reinforced lever press on both the first and last day of SA. No significant deviations in phasic firing were observed on the first day of SA (Figure 3C). In contrast, on the last day of SA, a significant increase in firing was observed immediately preceding and following the reinforced lever press specifically in the subpopulation of cells that was tonically inhibited during SA (Figure 3D).

**Figure 3.**
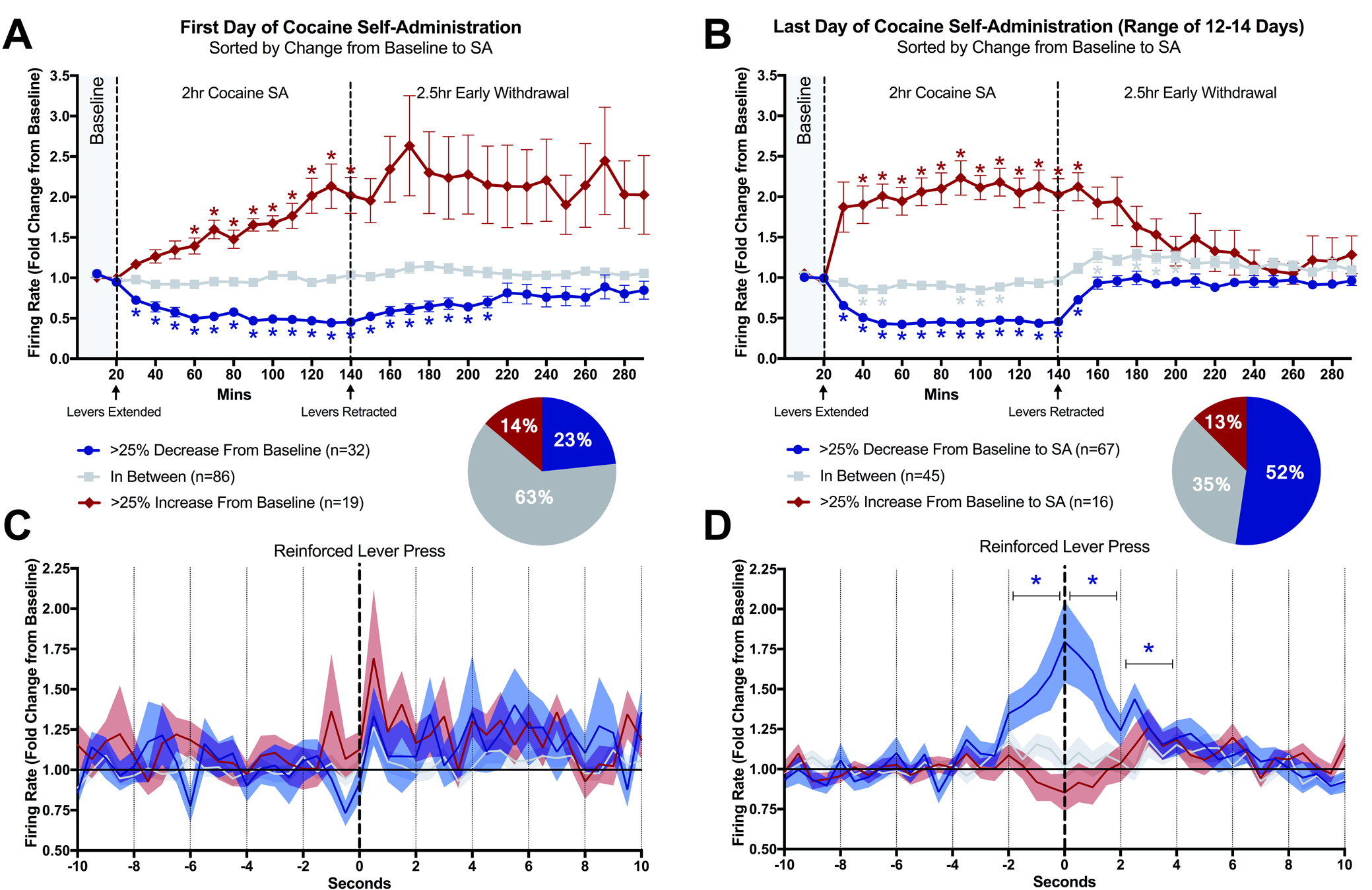
Representation of electrophysiological data from Experiment 1. Figures 3A and 3B show PL cortex activity before, during, and after cocaine SA for the first day and last day, respectively. Data are represented as average fold change (± SEM) from baseline and categorized into three different groups determined by the extent of variation from baseline during SA. Asterisks for figures 3A and 3B indicate a significant difference from baseline as indicated by a one-sampled t-test with Bonferroni correction for multiple comparisons (p<0.05). Figures 3C and 3D show spike firing rates in a time window from 10 sec before until 10 sec after a reinforced lever press for the first day and last day of SA, respectively. Data are represented as average fold change (± SEM) from baseline (the average firing rate of -15 to -10 sec) and categorized by the same criteria as figures 3A and 3B. For analysis, firing rate was compiled into 2-sec bins. Asterisks for figures 3C and 3D indicate a significant difference from baseline as indicated by a one-sampled t-test with Bonferroni correction for multiple comparisons (p<0.05).

### Experiment 2a: Delaying cocaine availability, but not operant responding or cue presentation, delays the tonic decrease in PL activity in rats during cocaine SA

We next tested whether the enhanced tonic inhibition of PL firing was caused by the cocaine itself or by the associated lever pressing and/or cues. Hence, in Experiment 2a, cocaine availability was delayed for the first 30 min of the self-administration session to determine whether operant responses in combination with cue presentation (in the absence of cocaine) were sufficient to reduce tonic PL cortical firing during cocaine SA. Repeated within-session one-sample t-tests (with Bonferroni correction) revealed that overall firing did not decrease until cocaine was made available (Figure 4A). This finding indicates that operant responding in combination with cue presentation in the absence of cocaine is not sufficient to alter the overall rate of firing.

**Figure 4.**
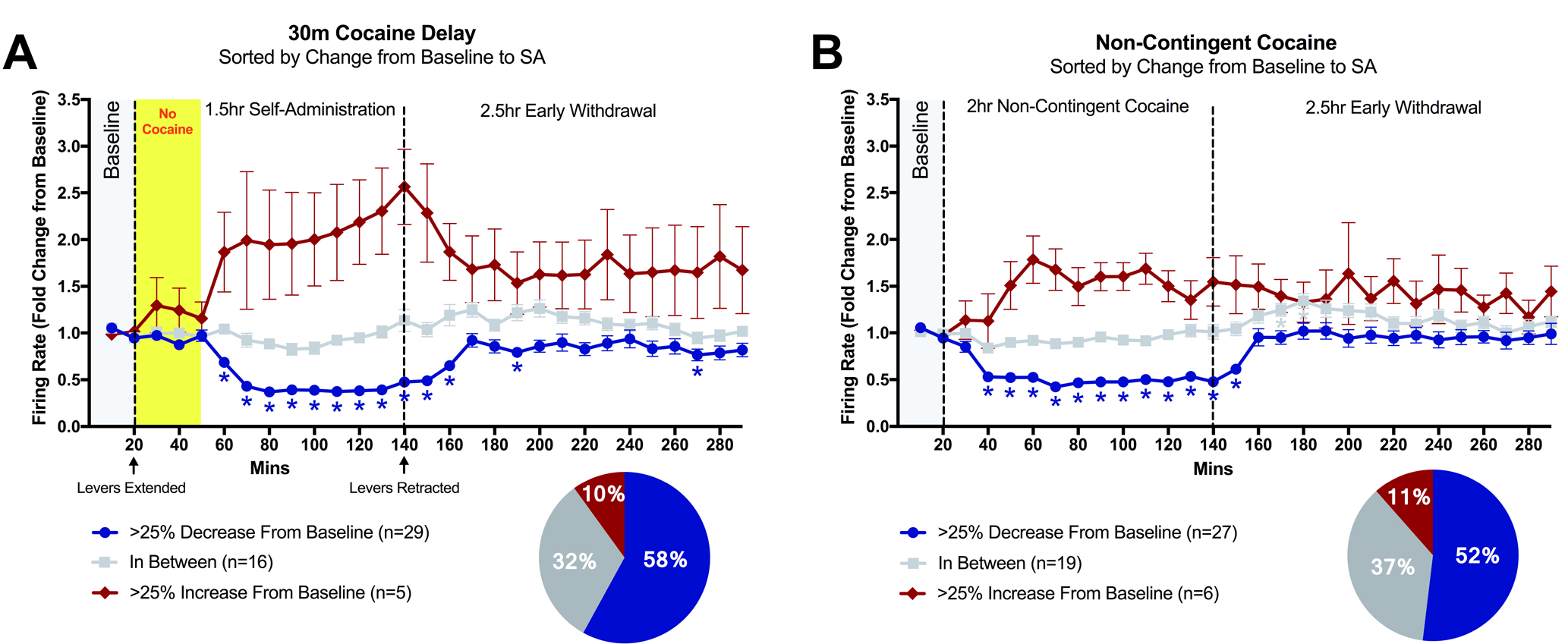
Representation of electrophysiological data from experiment 2. Figure 4A shows PL cortex activity during the baseline, the cocaine delay phase, cocaine SA, and early withdrawal for experiment 2a. Figure 4B shows PL cortex activity during baseline, non-contingent cocaine administration, and early withdrawal for experiment 2b. Data are represented as average fold change (± SEM) from baseline and categorized into three different groups determined by the extent of variation from baseline during cocaine administration. Asterisks indicate a significant difference from baseline as indicated by a one-sampled t-test with Bonferroni correction for multiple comparisons (p<0.05).

### Experiment 2b: Non-contingent cocaine decreases PL activity in rats with cocaine self-administration experience

Having shown that cocaine is necessary for PL suppression, we next tested whether cocaine by itself was sufficient for this effect in experiment 2b in which cocaine was non-contingently administered in the absence of operant responses (levers remained retracted) and cue presentations. Repeated within-session one-sample t-tests (with Bonferroni correction) revealed that, in a subset of neurons, firing is significantly reduced when compared to baseline (Figure 4B). These data indicate that cocaine by itself is sufficient to induce a decrease in tonic firing in the majority of cells in the absence of operant responding and drug-paired cues.

### Experiment 3: Yoked cocaine administered throughout the entire paradigm is not sufficient to produce a cocaine-induced transient tonic downshift in activity in the majority of PL cortex neurons

In Experiment 3, rats were yoked to the infusion schedule of self-administering rats. Cocaine was non-contingently administered and yoked rats received the identical cue-presentation procedures that self-administering rats received. Repeated within-session one-sample t-tests (with Bonferroni correction) revealed that subsets of cells were excited or suppressed during the SA phase of the paradigm on both the first and last day (Fig 5). However, a chi-square test revealed that the distribution of neurons that was excited/suppressed was not significantly changed between the first and last day of yoked cocaine (Chi-square: X^2^ (2, N =77) = 0.6882, p=0.71). This finding indicates that contingency of responding for cocaine during training is necessary to increase the number of cells that show a tonic suppression from the first to the last day.

**Figure 5.**
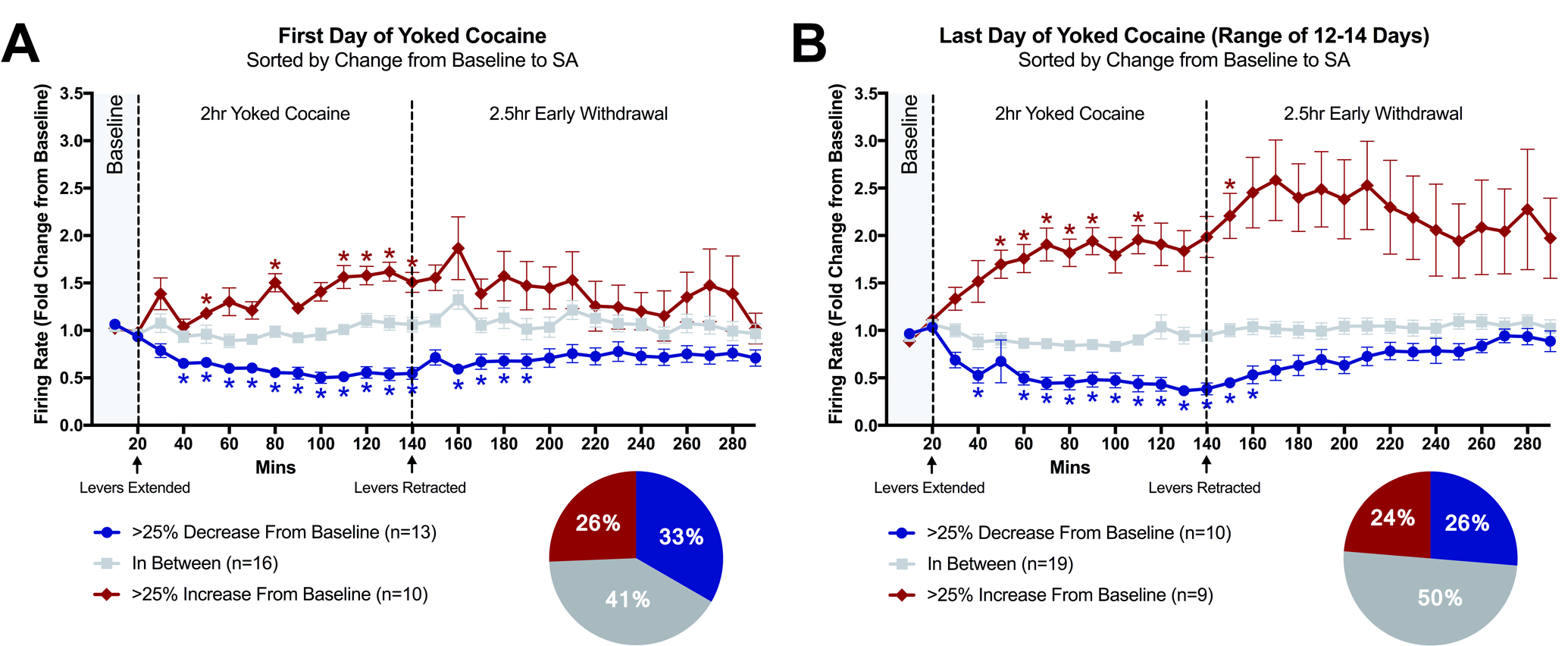
Representation of electrophysiological data from experiment 3. Figures 5A and 5B show PL cortex activity before, during, and after yoked cocaine for the first and last days, respectively. Data are represented as average fold change (± SEM) from baseline and categorized into three different groups determined by the extent of variation from baseline during yoked cocaine administration. Asterisks indicate a significant difference from baseline as indicated by a one-sampled t-test with Bonferroni correction for multiple comparisons (p<0.05).

As seen in Supplementary Figure 1*, a priori* two-tailed t-tests revealed a significant increase in the number of infusions from the first day to the last day in self-administering rats (t=3.741, df=34, p<0.05) and yoked cocaine rats (t=4.133, df=10, p<0.05). Importantly*, a priori* two-tailed t-tests indicated no difference on the first day between cocaine self-administering rats and yoked cocaine rats (t=0.4165, df=22, p=0.6811) and no difference on the last day between cocaine self-administering rats and yoked cocaine rats (t=1.058, df=22, p=0.3017). Taken together, these data suggest that infusion schedules for the yoked animals are representative of those for self-administering animals and indicates that the electrophysiological differences between these two groups are not due to variations in the number of infusions received.

## DISCUSSION

The data in this study show that after 12-14 days of SA, cocaine decreased overall tonic firing in the majority of recorded PL neurons. Moreover, there was a phasic increase in activity preceding and following the reinforced lever press specifically in those neurons that displayed a tonic decrease during cocaine SA. Investigations into the source of the downward tonic shift in activity revealed that cocaine was both necessary and sufficient for this effect to occur. In contrast, a yoked cocaine experiment revealed that non-contingent cocaine administered throughout the entire paradigm was not sufficient to induce this same tonic downshift in the majority of PL cortex neurons on the last day.

Chronic cocaine use has been shown to decrease baseline metabolic activity in the PFC of humans (Volkow *et al*, 1992), leading to the hypothesis that drug use over time contributes to a “hypofrontality” state (Bechara, 2005; Goldstein and Volkow, 2002). However, activity is increased in cocaine abusing individuals in response to drug-relevant information (Wang *et al*, 1999; Wilcox *et al*, 2011). Previous work in animals has shown similar alterations in the PL cortex with continued cocaine use. *Ex vivo* patch clamp recordings have shown decreased excitability in layer V pyramidal cells in rats that have undergone long-access cocaine SA (Chen *et al*, 2014). Also *in vivo* electrophysiological recordings in rats have shown decreased baseline activity with repeated cocaine SA (Sun and Rebec, 2006), yet enhanced firing in response to cocaine-related information (Sun and Rebec, 2006; West *et al*, 2014). All of these data point to the idea that there is a progressive dysfunction of prefrontal areas of the brain as cocaine use proceeds. It is likely that the tonic shift prompted by cocaine and the enhanced phasic responding surrounding drug-seeking in our current study reflect underlying maladaptive changes in the PL cortex induced by repeated cocaine exposure.

Of particular interest in the current study is the presence of a phasic increase in activity preceding and following the lever press specifically in those cells that display a tonic cocaine-induced decrease in activity. The implication of this phenomenon is that while downstream targets of the PL cortex receive less tonic input from the PL cortex during cocaine SA, this lower “floor” of activity may act to amplify the “signal-to-noise” for drug-related information coming from the PL cortex. There is evidence for cocaine SA-induced alterations in firing in downstream targets of the PL cortex. For example, cocaine-induced “session-long” decreases in activity have also been observed in the nucleus accumbens (primarily in the core subsection) of rats trained under a long-access cocaine self-administration paradigm (Peoples *et al*, 1998; 2004; 2007). Moreover, these cocaine-induced “session-long” decreases in accumbal firing show similarities to our current data, in that the extent of suppression is enhanced with continued SA experience when comparing day 15 to day 2 (Peoples *et al*, 1999). Because the nucleus accumbens core receives dense excitatory projections from the PL cortex, it is possible that these tonic decreases in the nucleus accumbens are the result of upstream tonic decreases in the PL cortex (although a “local” effect of cocaine in the nucleus accumbens core cannot be ruled out). Together, these overall decreases provide lower background activity, which may allow for the amplification of drug-relevant information and the focusing of drug-seeking behavior while cocaine is in the system. Future studies should establish pathway specificity to determine if PL neurons that display a cocaine-induced tonic decrease in activity project preferentially to the accumbens core or to another area of the brain.

Based on waveform and firing rate criteria, it is likely that most of our recorded neurons are pyramidal neurons (>0.5ms peak-to-valley waveform and <10Hz), and not GABAergic interneurons, which tend to have shorter duration action potentials and higher firing rates. It is possible that the cocaine-induced reductions in activity in our experiments are the result of increased dopamine activation of interneurons in the PL cortex. Cocaine exerts its pharmacologic effect primarily by blocking dopamine transporters (Sulzer and Sulzer, 2011), thereby prolonging stimulation of dopamine receptors. To this point, cocaine SA significantly increased extracellular dopamine in the medial prefrontal cortex of rats after 1, 8, and 17 one hr sessions of cocaine self-administration with the 17^th^ session yielding the greatest increase in dopamine levels (Ben-Shahar *et al*, 2012). Activation of dopamine receptors increases extracellular GABA levels in the medial prefrontal cortex of both naïve (Grobin and Deutch, 1998) and cocaine-sensitized (Jayaram and Steketee, 2005) rats. The ability of dopamine to modulate pyramidal neuron activity depends on the type of dopamine receptor activated and the type of cell on which these receptors are expressed. Direct D2 receptor stimulation of layer V pyramidal neurons decreases their activity *in vivo* (Godbout *et al*, 1991; Pirot *et al*, 1992). Further, stimulation of the VTA to induce dopamine release increases firing in prefrontal inhibitory fast-spiking interneurons and decreases firing in pyramidal cells paired with these interneurons (Tseng *et al*, 2006). D2 receptors appear to mediate these effects in mature rat prefrontal cortex *in vitro* (Tseng and O'Donnell, 2007). If D2 receptors on fast-spiking interneurons are sensitized with continued cocaine use in our current set of studies, increases in dopamine induced by cocaine would activate these inhibitory cells to clamp down activity of pyramidal cells. Future studies are needed to investigate the underlying mechanisms of cocaine-induced changes in neuronal activity by examining the effects of local PL infusion of D1 and D2 receptor antagonists on the cocaine-induced decrease in tonic activity.

While our work expands upon previous *in* vivo electrophysiological studies, there are certain key findings that our data do not replicate. Sun and Rebec (2006) described decreased baseline firing rates with continued cocaine SA experience. In contrast, we did not find a statistically significant decrease in baseline firing from the first SA day to the last SA day (data not shown). This discrepancy is possibly due to differences in the way that each paradigm was designed. Notable differences include the schedule of reinforcement (FR 1 vs. FR 5), operant pretraining (no pretraining vs. sucrose pretraining), baseline measurements (20 min vs. 1 min), and dose of cocaine used (0.2mg/50µl/infusion vs. 0.125mg/50µl/infusion).

Taken together, our data describe a novel phenomenon by which cocaine SA induces large tonic decreases and phasic increases in PL cortical activity in a time-dependent manner. Further, contingency during training is necessary to induce changes in the PL cortex that prime this area of the brain to decrease its overall activity when cocaine is on board and suggest that these repeated, daily decreases in activity that accumulate over time may lead to dysfunctions in excitation-transcription coupling that underlie cocaine-seeking (Go *et al*, 2016; Whitfield *et al*, 2011). Future work will focus on determining the neural mechanisms involved in this effect, including the identification of specific neurotransmitter systems, determination of the cell subtype of recorded neurons, and the identification of patterns of firing in neurons projecting to specific downstream targets of interest.

## Acknowledgements

We would like to thank Ben Siemsen, Erica Herzig, Jordan Hopkins, and Dr. Casey O’Neill for performing catheter surgeries. We also thank Nicki Pullmann for her assistance in the preparation of brain tissue for histology. Lastly, we thank Jasper Heinsbroek for his generous contribution of Matlab code that greatly accelerated data analysis.

